# KISS1 is a potential immunological and prognostic biomarker for cancer metastasis in pan-cancer and affects breast cancer in a different way

**DOI:** 10.1101/2023.07.05.547825

**Authors:** Hao Jiang, Jianru Xiao, Hao Zhang, Bo Li, Wei Xu, Zhenxi Li

## Abstract

Metastasis to other organs accounts for the majority of death in cancer patients. Metastatic capacity enables tumor cells resist to chemotherapy. KISS1 had been found to suppress metastasis in cancers like melanoma, but improve the metastatic phenotype of breast cancers, which means its role in tumor metastasis is tissue context-dependent. This character suggests the KISS1 might have promising potential to interpret tumor metastasis in a pan-cancer perspective. Here, we analyzed the potential functions of KISS1 in pan-cancer, especially in breast cancer, by the datasets of TCGA and GTEx. KISS1 exhibited a different expression status between mRNA level and protein level in pan-cancers, with a significant association between KISS1 expression and prognosis of cancer patients. Besides, the correlation between KISS1 expression and immune cell infiltration, immunosuppressive cells, immune checkpoint genes, TMB, MSI, tumor purity and HRD indicates the potential role in regulating immunotherapy. Thereafter, we further clarified the role of KISS1 in promoting metastasis in breast cancer. Furthermore, we found that the post-translational modification may explains the KISS1 protein stability in breast cancers. In summary, our first pan-cancer study offers new insights and relatively comprehensive understandings of the oncogenic roles of KISS1 across various cancers.

## Introduction

The durable control of cancers remains to be a challenge for medical workers, especially when metastases develop, though there has undergone a revolution in cancer treatment and early detection [1–3]. Metastasis to other organs, even a preference for some specific organs [4], represents a major obstacle towards cancer therapies [5]. The drug resistance of tumor cells accounts for the major reason which leads to an unsatisfactory outcome for neoplastic diseases treatment [6], and the drug resistance is associated to the promotion of metastatic ability with the occurrence of epithelial to mesenchymal transition (EMT) [7–8]. The development and outgrowth of cancer metastasis depend on several positive and negative regulators [9], including metastasis suppressor gene or protein that subjected to modifications during gene transcription and protein translation [10]. Thus, several metastasis suppressors have been described [11], and among them KISS1 raised our interesting for the still controversial role in tumor metastasis [12].

Physiologically, KISS1 could binding to G protein-coupled receptor (GPR54, also named KISS1R) to activating hypothalamic-pituitary-gonadal axis, which regulates puberty and reproductive functions [13]. Originally, its metastasis suppressor role was identified in melanoma cells, in which cancer metastasis had been suppressed by introducing a human chromosome 6[14]. Concordantly, other researches have confirmed the metastasis-suppressive role of KISS1 in lung, prostate and pancreatic, mainly due to the KISS1 receptor and the negative regulation of CXCR4 (CXC chemokine receptor 4) [15–17]. However, KISS1 had been reported to promote the metastatic capacity of estrogen-receptor (ER) negative mammary epithelial and breast cancer cells [18]. Therefore, the role in cancer metastasis of KISS1 is context-dependent, which needs further studies to make out the underlying mechanism.

Given all the previous studies, there is no studies of KISS1 in an entire cancer spectrum. Here, we did this pan-cancer study by public databases to acquire a most efficient understanding of KISS1 in terms of molecular mechanisms and predictive value in tumor biology [19–20]. Technologically, we included a number of factors, such as gene expression, protein expression, survival status, immune infiltration, immunosuppressive cells, tumor mutational burden (TMB), tumor mutational burden (MSI), rumor purity, (Homologous recombination deficiency) HRD, cell experiments and post-translational modification to investigate the potential molecular mechanism of KISS1 in the pathogenesis and clinical prognosis of various cancers.

## Materials and methods

### mRNA and protein expression analysis

For the mRNA expression, we downloaded 10535 pan-cancer samples from TCGA and 15776 data combined with the GTEx (https://commonfund.nih.gov/GTEx/)) resource from the UCSC (https://xenabrowser.net/) database. The mRNA sequencing data for KISS1 in 34 tumors and 31 normal tissues was transformed by log2(x+0.001) [21–24]. The protein expression analysis of CPTAC ((Clinical proteomic tumor analysis consortium) dataset was conducted on UALCAN portal ((http://ualcan.path.uab.edu/analysis-prot.html)[25–26]. Besides, we explored the protein expression statues by immunohistochemical samples from HPA (Human proteins atlas) web (https://www.proteinatlas.org/) [27–28].

### Correlations between KISS1 expression and clinical features

The analysis of relationship between clinical stage, metastasis statues and KISS1 expression was done by R software (version3.6.4) and obtained violin plots of KISS1 expression in different pathological stage of some indicated tumors by the GEPIA2 (Gene expression profiling interactive analysis, version 2) web server (http://gepia2.cancer-pku.cn/#analysis) [29].

### Survival prognosis assessment

The prognostic value of KISS1 in various cancers was assessed by 2 types of clinical outcomes: overall survival (OS), disease-free survival (DFS). High-quality prognostic expression data were picked up from the pan-cancer expression data, and the results were visualized using “ Quick select” section on cBioPortal web (https://www.cbioportal.org/) [30–31]

### Relationship analysis of immune traits

We explored the association between KISS1 and immune infiltrates in pan-cancer from TCGA by the “Immune-Gene” module of the TIMER2 web server (http://timer.cistrome.org/)[32]. The StromalScore, ImmuneScore, and ESTIMATE scores were obtained by the R package “ESTIMATE” for all the tumor samples [33]. KISS1 and 8 immune checkpoint pathway genes were collected from a pan-cancer gene expression dataset [34]. As for the relationship with immunosuppressive cells, we selected myeloid-derived suppressor cells (MDSCs) and regulatory T cells for the analysis [34]. The CIBERSORT, CIBERSORT-ABS, XCELL and EPIC algorithms were applied for immune infiltration estimations [26]. We preformed the purity-adjusted Spearman’s rank correlation text to get the P-values and partial correlation values. These data were demonstrated as heatmap and a scatter plot.

### Relationship between KISS1 and genomic heterogeneity and stemness

Analysis of genomic heterogeneity of tumors, including TMB, MSI, purity, HRD, was performed to predicts the response to immune checkpoint inhibitors (ICIS) [35–37]. Standardized pan-cancer dataset was downloaded from UCSC (https://xenabrowser.net/) and GDC(https://portal.gdc.cancer.gov/), and TMB score was acquired by the R package “maftools”. MSI scores, purity and HRD data were acquired from the research of Thorsson V et al and the spearman correlation analysis was performed [23].

### Cell culture, transfections, immunoblotting and immunoprecipitation

MBA-MD-231 was purchased from ATCC and the highly bone-metastatic (BM) derivatives were obtained from Dr. Zhenxi LI (Changzheng Hospital, 415 Feng Yang Road, Shanghai 200003, People’s Republic of China). All the cells were cultured in DMEM (BasalMedia, L110KJ), with 10% FBS (YLESA, S211201) and 1% 100x Penicillin-Streptomycin solution (YEASEN, 60162ES76), and grown at 37°C in a 5% CO2 incubator. Human KISS1 was cloned into pLVX-neo vector (Clontech) for overexpression, and the annealed sense and antisense shRNA oligonucleotides were cloned into pLKO.1-puro vector (Addgene) for knockdown of human KISS1. The shRNAs used were: TRCN0000059065 (#1), TRCN0000059064 (#2), TRCN0000059067 (#3). Flag-WWP1 was transfected into MDA-MB-231 by PEI (Polysciences, 24765-100) according to the manufacturer’s protocol. During immunoprecipitation assays, cells was lysed with RIPA (Beyotime, P0013D) containing protease inhibitors (Sigma), and the immunoprecipitations were performed using anti-Flag M2 beads (Sigma, M8823) at 4°. The immunoprecipitates was washed with NETN buffer. Immunoblot (IB) analysis was performed using specific antibodies and related secondary antibody and the visualization was obtained with chemiluminescence. Antibodies used in IB analysis: anti-KISS1 (Sigma-Aldrich, MABC60), anti-GAPDH (Abcam, ab8245), anti-Flag (Abcam, ab205606).

### Cell colony formation and transwell assay

Cells were positioned on a six-well plate and cultured for 2 weeks, during which time the medium was changed every 3 days. Formed colonies were fixed in 4% Paraformaldehyde for 30mins and stained with Crystalline Violet (Sigma-Aldrich, C0775) for 10mins, and photographed and analyzed by ImageJ software. As for the transwell assay, 8000 cells were inoculated in each tranwell chambers (Corning, CLS3378), with 200ul DMEM containing 1% FBS, and 600ul culture medium containing 20% FBS was added in the lower chamber.

### Statistical analysis

Gene expression differences was compared by the Kruskal-Wallis and the Wilcoxon Rank Sum Test. Correlations between the two groups were calculated by Spearman or Pearson correlation analysis. Survival curves were generated by Kaplan-Meier method and Cox regression analysis. Categorical variables were analyzed using Chi-square test and Fisher’s exact test. Statistical significance was generated using log-rank test and the level of significance was set at P < 0.05 (*), P <0.01 (**) and P < 0.001 (***). The above visualization was performed with R software (version 3.6.4), GraphPad Prism 9 and Sangerbox web (http://www.sangerbox.com/) [38].

## Results

### Gene expression of KISS1 in pan-cancer

To obtain the expression status of KISS1 in pan-cancer, we analyzed 33 cancers based on TCGA databases. The KISS1 was significantly upregulated in 13 cancers, including LUAD, COAD, COADREAD, BRCA, STES, STAD, UCEC, LIHC, THCA, READ, PAAD, BLCA and CHOL (Figure 1A). However, we also find that the KISS1 was downregulated in KIRP, KIPAN and LUSC (Figure 1A). Furthermore, we explored the database from GTEx to evaluate the difference of KISS1 expression between normal tissues and tumor tissues. As showed in Figure 1B, the KISS1 was upregulated in 19 cancers, such as UCEC, BRCA, CESC, LUAD, ESCA, STES, COAD, COADREAD, STAD, LIHC, BLCA, THCA, READ, OV, PAAD, UCS, ALL, ACC and CHOL, and downregulated in 13 cancers, including GBM, GBMLGG, LGG, KIRP, KIPAN, PRAD, KIRC, LUSC, WT, SKCM, TGCT, LAML, KICH (Figure 1B).

**FIGURE 1.**
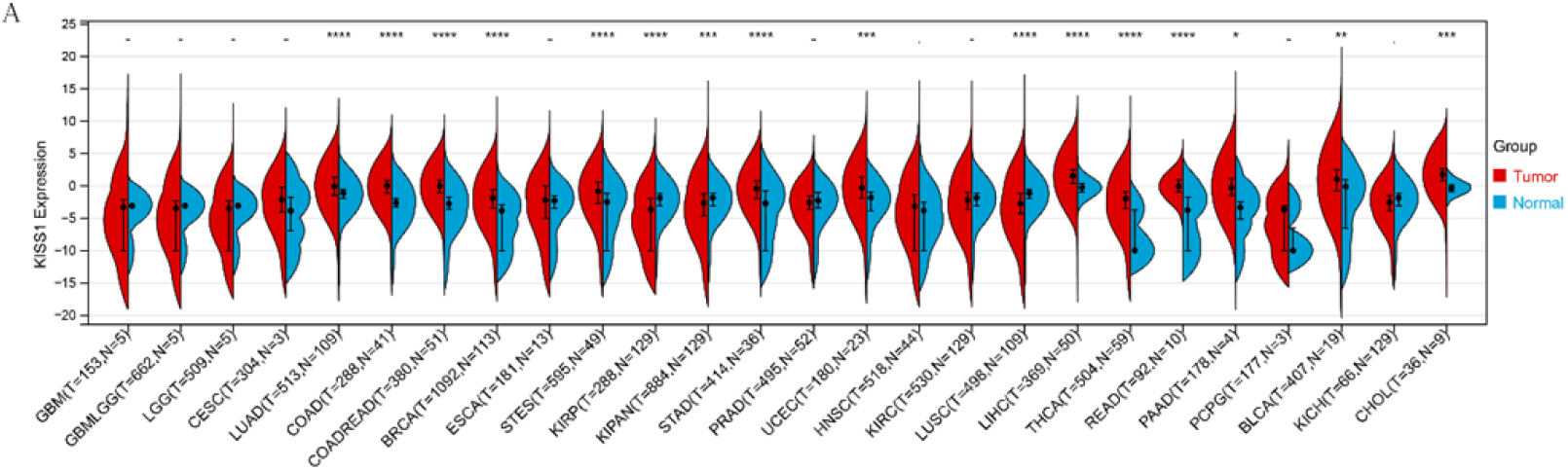
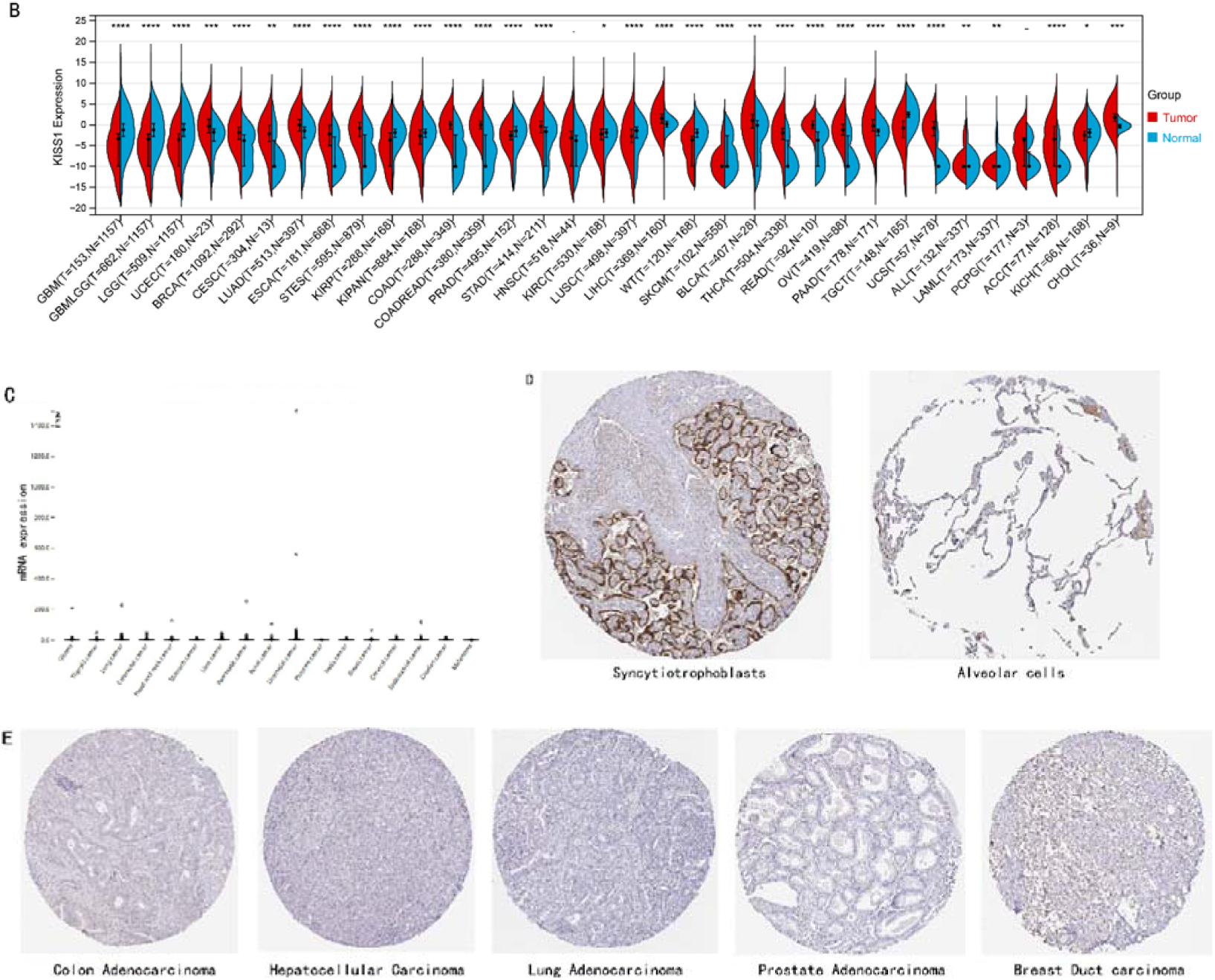
Expression level of KISS1 mRNA and protein in pan-cancer. (A) The expression status of KISS1 gene in tumor tissues by data from the TCGA database. (B) Analysis of KISS1 gene expression between tumor tissues from TCGA and corresponding normal tissues from GTEx database. Ns, p≥0.05; *p<0.05; **p<0.01; ***p<0.001; ****p<0.0001. (C) mRNA expression level of KISS1 in pan-cancer from TCGA database. (D) The KISS1 expression of human normal tissues in immunohistochemical from HPA dataset. (E) The protein expression of KISS1 in human cancer tissues from HPA dataset.

### Protein expression of KISS1 in pan-cancer

These results made us interested in the protein expression status of KISS1 in various cancers, unfortunately there is no public database available from CPTAC (Clinical proteomic tumor analysis consortium) dataset [25], which indicating the recognition to KISS1 remains superficial. By exploring the database from HPA (Human protein atlas), we found that KISS1 had a high mRNA expression level in the Urothelial cancer, and a relatively high level in Glioma, Lung cancer, Pancreatic cancer (Figure 1C). In terms of the protein expression level in human tissues, the KISS1 could physiologically detected in tissues like Syncytiotrophoblasts (HPA035542), mainly located in Cytoplasmic and membranous within cells, but nearly no expression in other issues like lung (CAB017775) (Figure 1D). Surprisingly, according to the databases in HPA, except in breast duct carcinoma (CAB017775), no KISS1 protein was detected in cancers like Colon Adenocarcinoma (HPA035542), Hepatocellular Carcinoma (HPA035542), Lung Adenocarcinoma (HPA035542) and Prostate Adenocarcinoma (HPA035542) (Figure 1E). The contradictory results between mRNA and protein expression in pan-cancers call for further investigations of KISS1, especially in terms of the post-translationally modification.

### Expression analysis of KISS1 in different pathological and clinical stage

In a prospect view of pan-cancer, we assessed the importance of KISS1 expression in different clinical and pathological stage. In COAD, COADREAD, BRCA, KIRP, KIPAN and KIRC the KISS1 demonstrates a differential expression status in different clinical and pathological stage (Figure 2A), and the detail information of COAD, COADREAD, BRCA, KIRP, KIPAN, KIRC were showed in Figure 2B. Besides, as the KISS1 was reported as a metastasis suppressor [12], we investigated the expression differences between different metastasis status in pan-cancers. Interestingly, according to datasets from TCGA, only in ESCA, KIPAN, PRAD, KIRC, PAAD, TGCT, SKCM it makes sense (Figure 2C). However, KISS1 was reported to suppress various cancers metastasis, such as melanoma, bladder cancer and lung cancer [12–15]. The contradictions indicated that the analysis of KISS1 is underinvested, especially in terms of its prognostic value.

**FIGURE 2.**
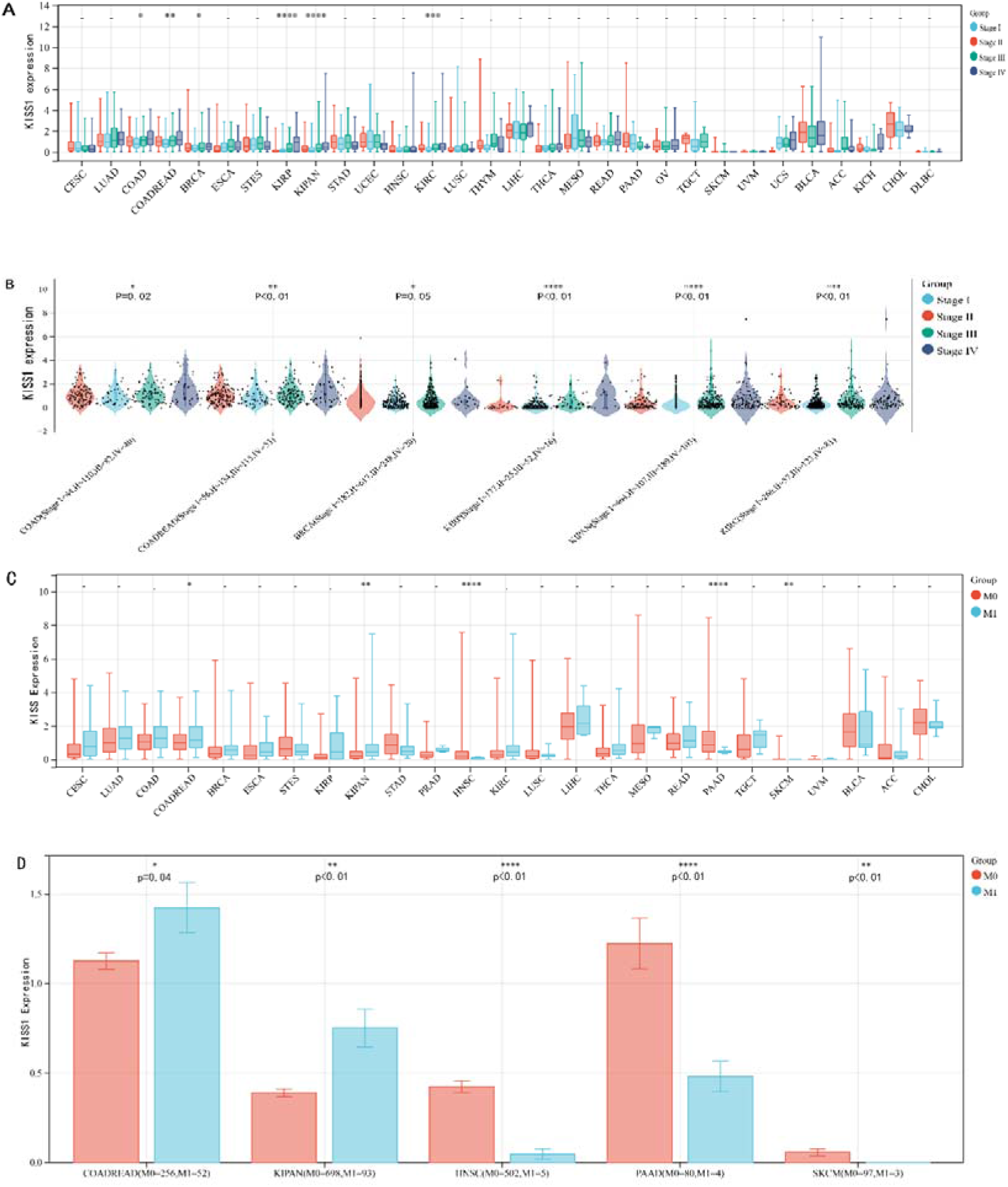
Expression analysis of KISS1 in different pathological and clinical stage. (A-B) KISS1 expression at different clinical stages; (C-D) KISS1 expression at different metastasis statues.

### Survival analysis

For the prognostic value, we first adopted cox regression model and Kaplan-Meier (KM) survival analysis of KISS1 in OS (overall survival) and DFS (Disease-free survival) [26]. According to the expression levels of KISS1, all the cancer cases in TCGA and GEO were divided into two groups, high-expression and low-expression groups. Then we investigated the correlation of KISS1 expression with the prognosis of patients from these datasets. Within the TCGA project, high expression of KISS1 contributes to a poor prognosis of OS for cancers of KIRP, KIRC, LUAD. While, the poor prognosis of OS in BLCA are related to the low expression of KISS1 (Figure 3A). Besides, high expression of KISS1 was a risk factor for DFS in patients with CESC, COAD, KIRP, PAAD (Figure 3B). And all these relationships between KISS1 and OS, DFS were further demonstrated by Kaplan-Meier (KM) survival analysis curves (Figure 3A and 3B). By comparison, the KISS1 expression correlated both with OS and DFS in KIRP.

**FIGURE 3.**
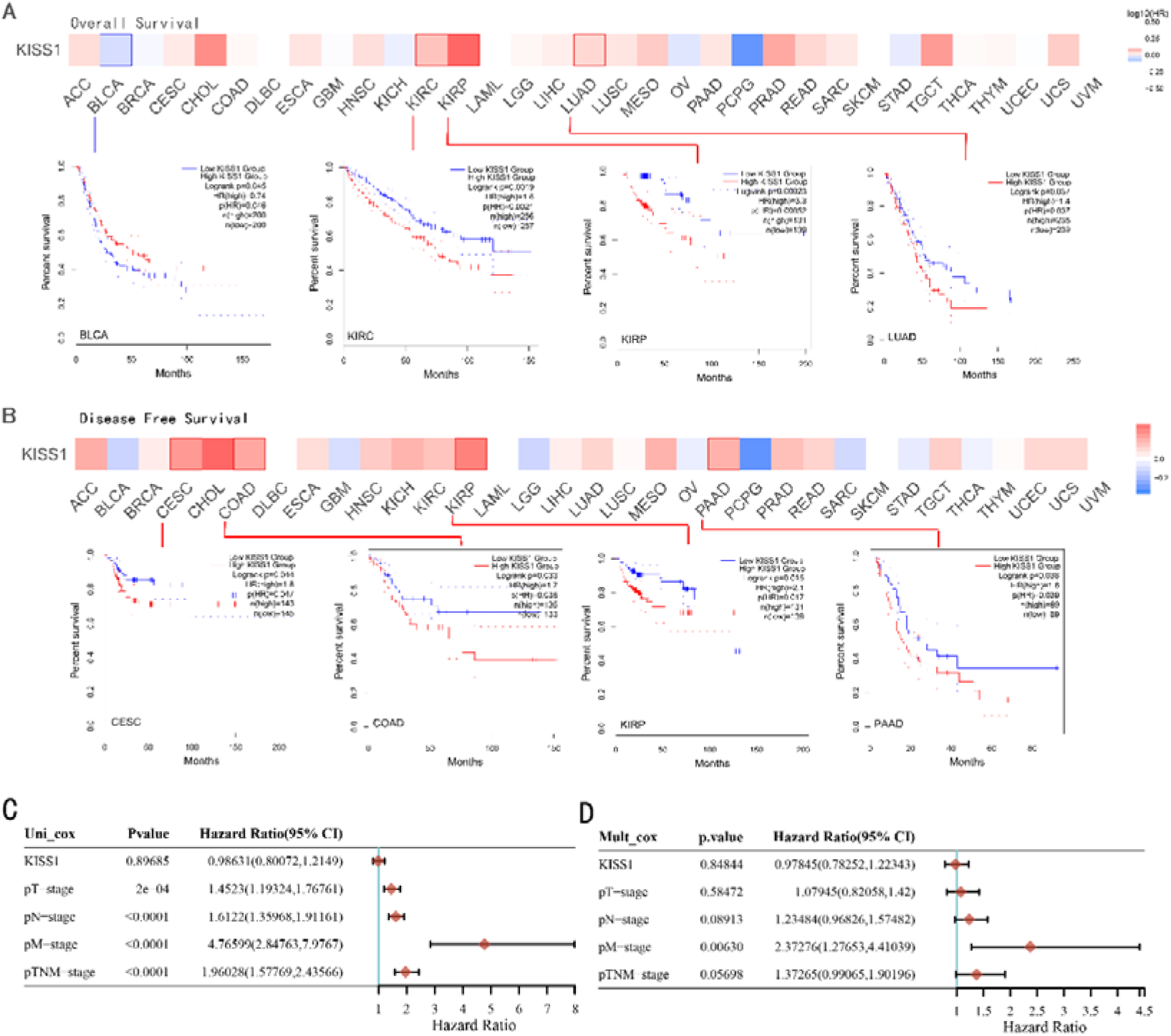
Relationship between KISS1 expression and survival prognosis of pan-cancers in TCGA. (A) Survival map and Kaplan-Meier curves of OS (Overall survival) of various tumors by KISS1 gene expression; (B) Survival map and Kaplan-Meier curves of DFS (Disease free survival) of various tumors by KISS1 gene expression; (C-D) Prognostic significance of KISS1 in breast cancer by univariate and multifactorial COX analysis.

### KISS1 is significantly related to immune infiltration

To investigating the potential role of KISS1 in the tumor microenvironment (TME), we explored the relationship between KISS1 and the level of immune infiltration in various cancers from TCGA by three immune scores and we picked up the four most significant ones respectively. Firstly, according to the StromalScore[33], KISS1 expression in STAD (N=388,R=-0.25,P=6.8e-7), BRCA (N=1077,R=-0.13,P=2.6e-5) and LIHC (N=363,R=-0.25,P=1.5e-6) was prominently negative correlated with immune infiltration (Figure 4A).While in HNSC (N=517,R=0.15,P=5.6e-4), the KISS1 expression was positive correlated with the immune infiltration (Figure 4A). Then as for the ImmuneScore [33], the expression of KISS1 was significantly negatively correlated with immune infiltration in COAD (N=282, R=-0.20, P=6.1e-4), COADREAD (N=373, R=-0.21, P=4.2e-5) and STAD (N=388, R=-0.30, P=1.4e-9), and positive correlated with immune infiltration in PRAD (N=495, R=0.20, P=5.1e-6) (Figure 4B). Lastly, by the EstimateScore, KISS1 expression in SARC (N=258, R=-0.22, P=4.1e-4), STAD (N=388, R=-0.30, P=1.4e-9) and LIHC (N=363, R=-0.20, P=1.4e-4) was negative correlated with immune infiltration (Figure 4C). While, there was a positive correlation between KISS1 expression and immune infiltration in PRAD (N=495, R=0.19, P=2.6e-5) (Figure 4C). In summary, we could find a mostly consistently trend that KISS1 negatively regulates TME in most cancers from TCGA, and only in cancers like PRAD the KISS1 expression and immune infiltration were both highly. However, the relationship between KISS1 expression and immune infiltration need further investigations before we could make a clear conclusion.

**FIGURE 4.**
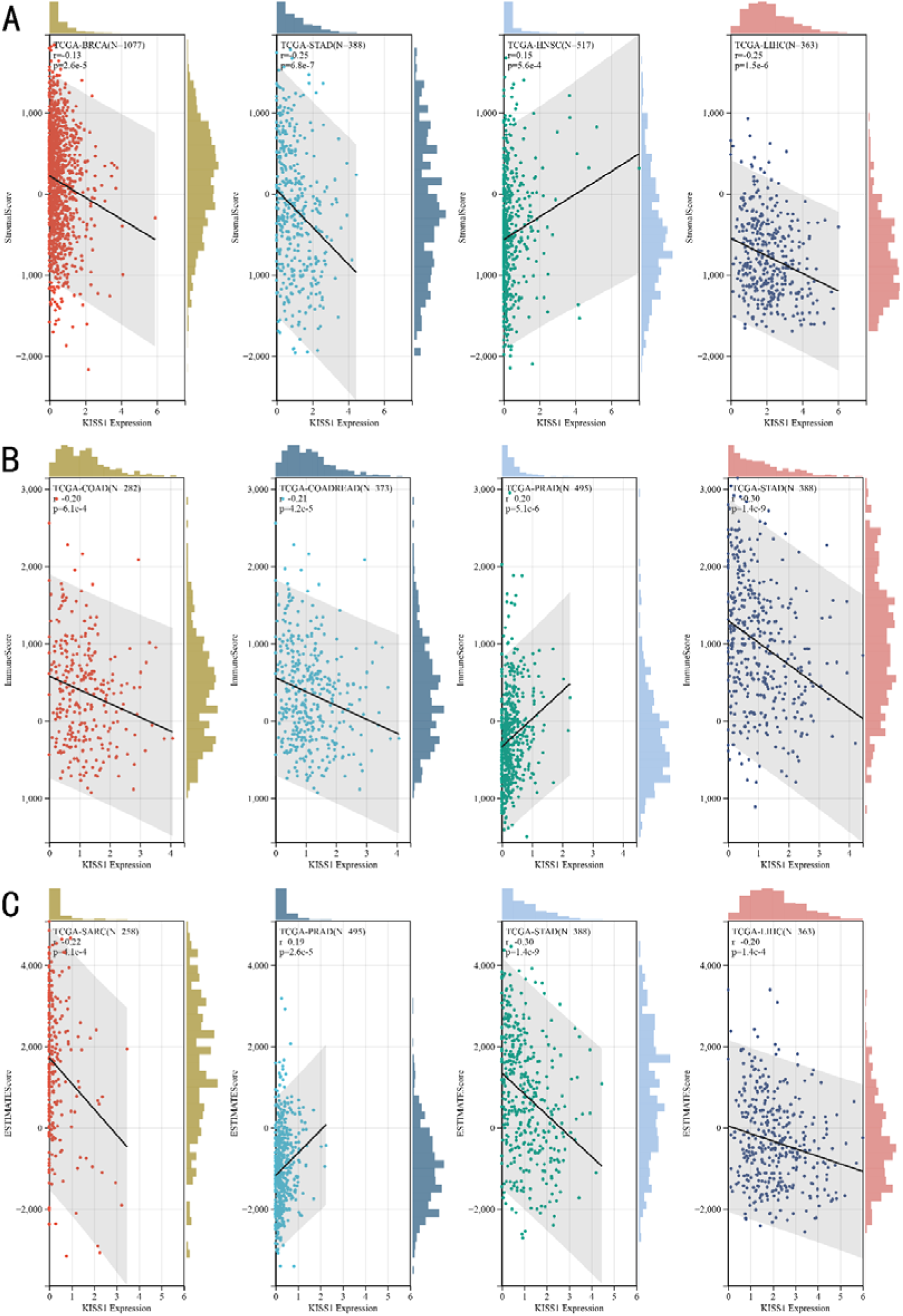
Relationship between KISS1 expression and immune infiltration. (A) Correlation of KISS1 expression with StromalScore; (B) Correlation of KISS1 expression with ImmuneScore; (C) Correlation of KISS1 expression with EstimateScore.

### KISS1 interacts evidently with immunosuppressive cells

Previously, we found that the KISS1 expression may negatively regulates TME. Furthermore, we exploring the importance of KISS1 on the suppressive tumor microenvironment (STM), through the relationship between KISS1 expression and the infiltration level of myeloid-derived suppressor cells (MDSCs) and regulatory T cells (Tregs) [39]. As a result, we observed a statistical positive correlation of KISS1 expression and immune infiltration of MDSCs in tumors of BRCA, BRCA-Basal, BRCA-LumA, BRCA-LumB, CESC, KIRC, KIRP, LGG, LIHC, MESO, PAAD, SARC, SKCM, SKCM-Metastasis, STAD, TGCT and UVM, but also a negative correlation for tumors only in BLCA and ESCA (Figure 5A), based on TIED algorithms. And the scatterplot data of BLCA, ESCA, BRCA, BRCA-Basal, BRCA-LumA, BRCA-LumB, PAAD, STAD were listed in Figure 5B.

**FIGURE 5.**
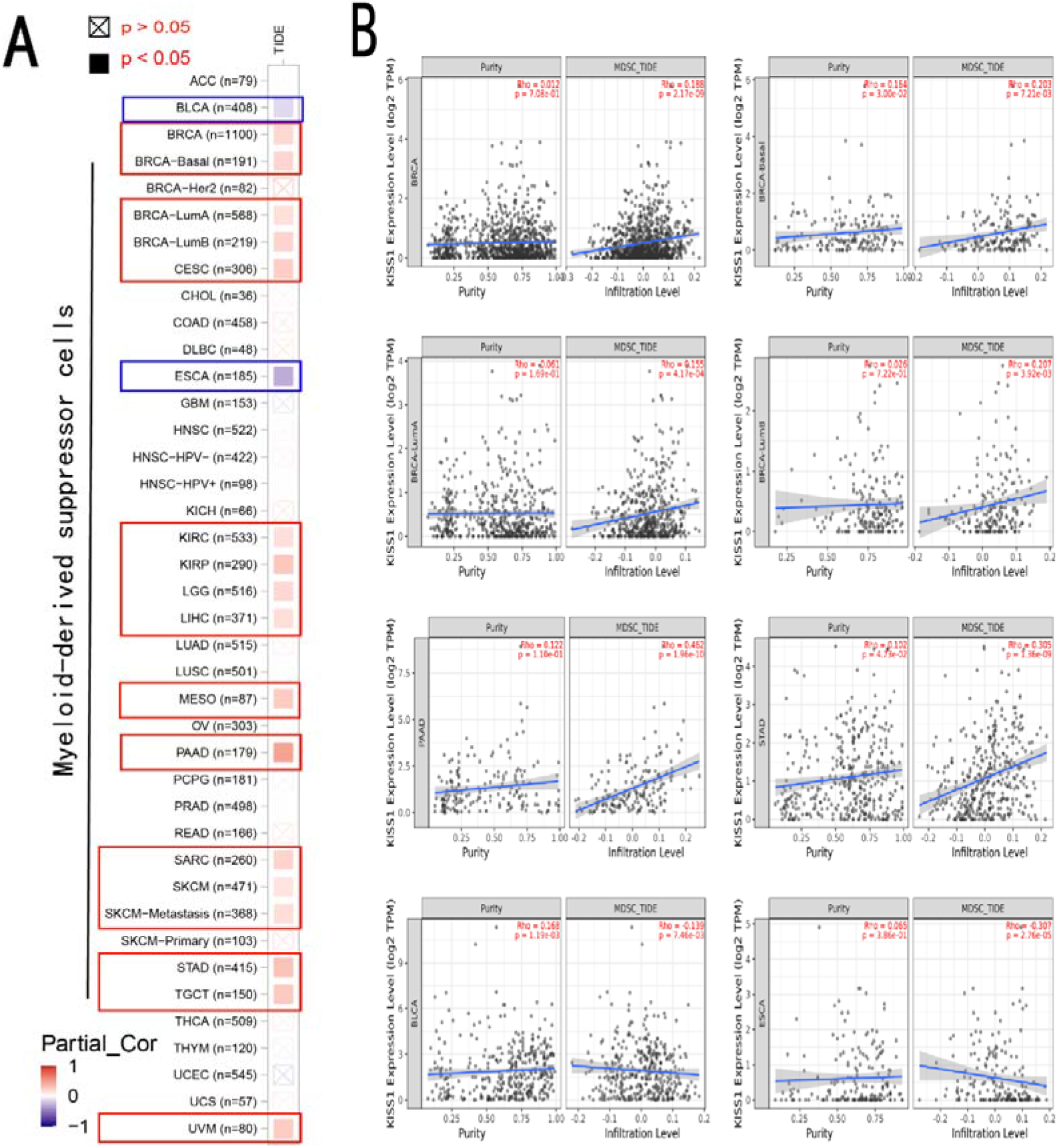
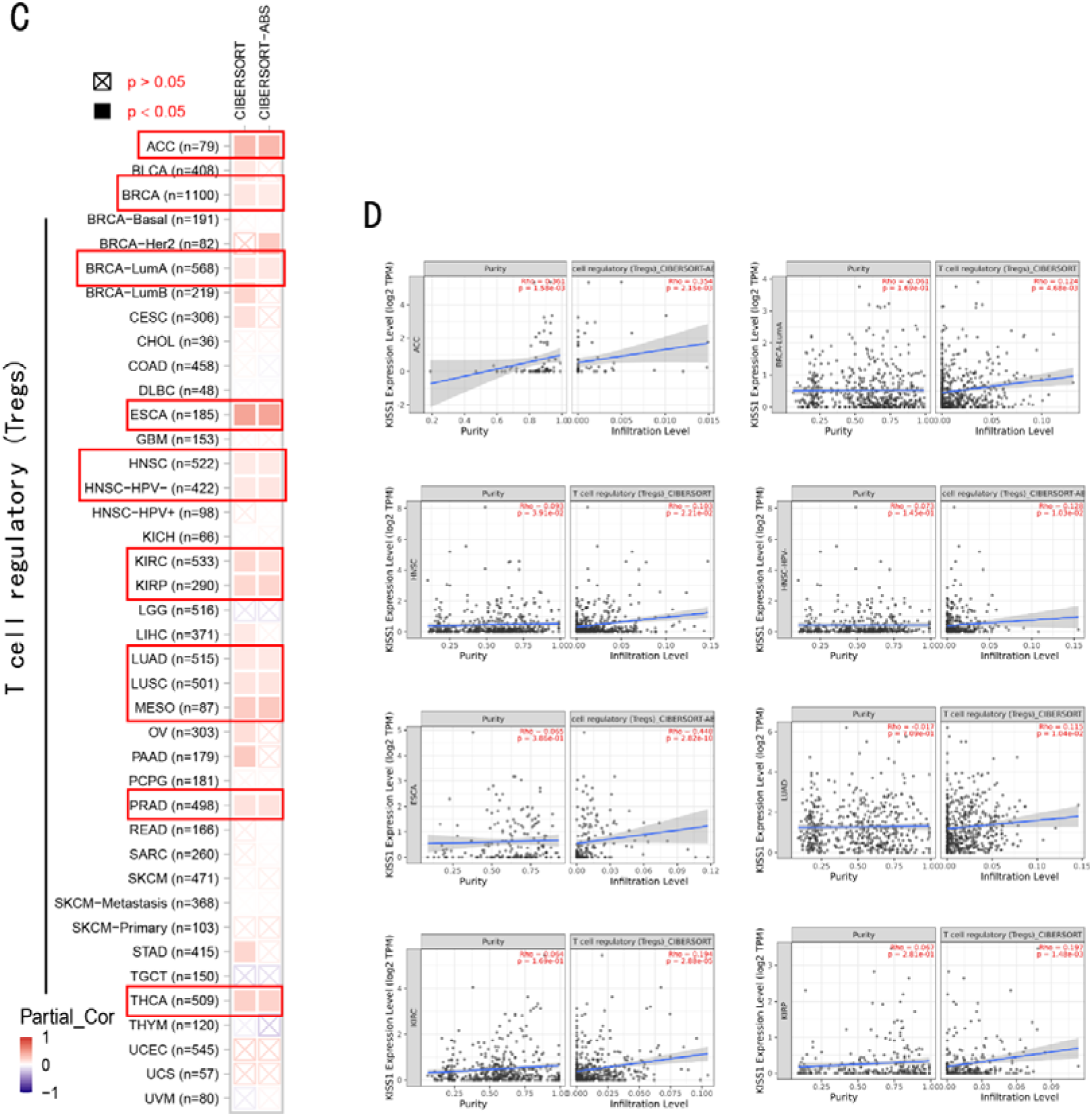
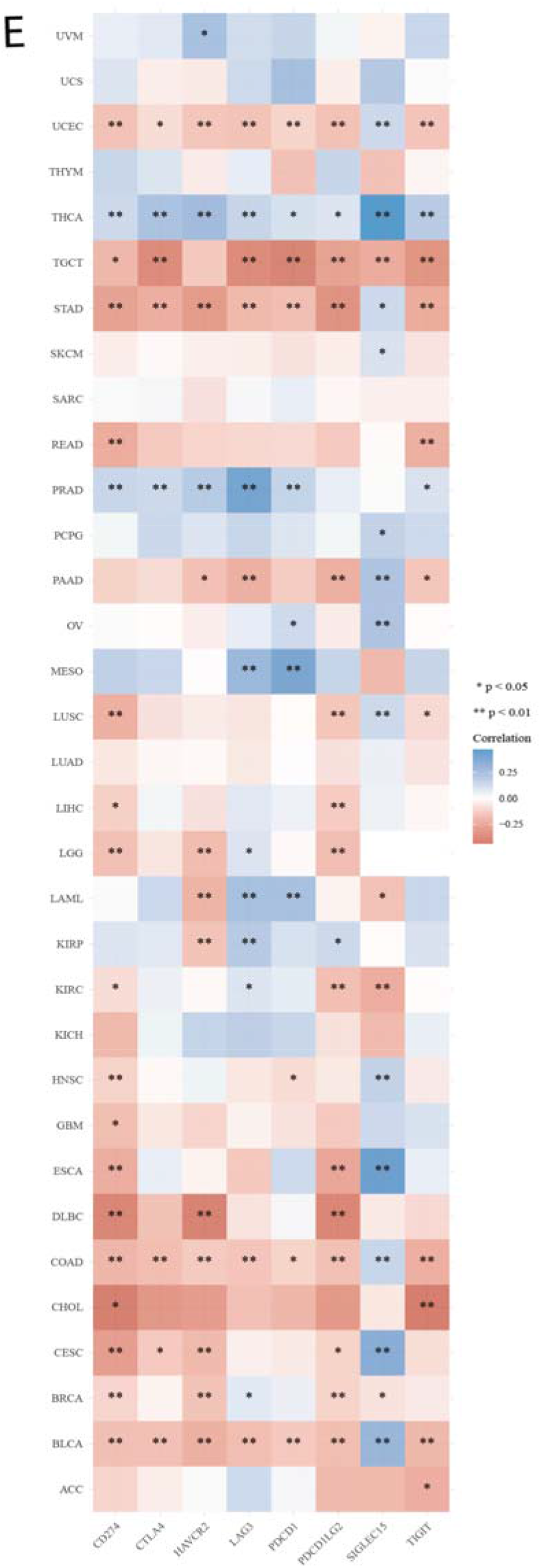
Correlation analysis between KISS1 expression and immunosuppressive cells infiltration. (A-B) Correlation between KISS1 expression and myeloid-derived suppressor cells (MDSCs) infiltration; (C-D) Correlation between KISS1 expression and regulatory T cells (Tregs) infiltration; (E) Relationship between KISS1 and chemokine genes.

Followed by MDSCs, we then analyzed the relationship between KISS1 expression and regulatory T cells (Tregs) infiltration. According to the CIBERSORT and CIBERSORT-ABS algorithm, we find a statistical positive correlation between the immune infiltration of Tregs and KISS1 expression in tumors of ACC, BRCA, BRCA-LumA, ESCA, HNSC, HNSC [HPV (human papillomavirus)-], KIRC, KIRP, LUAD, LUSC, MESO and THCA (Figure 5C), and the scatterplot data of ACC, BRCA-LumA, HNSC, HNSC [HPV (human papillomavirus)-], KIRC, KIRP, ESCA and LUAD were showed in Figure 5D. Interestingly, there is no negative correlation between Tregs infiltration and KISS1 expression in pan-cancers according to the CIBERSORT and CIBERSORT-ABS algorithm.

Indeed, the infiltration of immunosuppressive cells could be a major reason for undermining the therapeutic efficacy of ICIs [40]. Therefore, we further analysis the correlation of KISS1 expression with 8 immune checkpoint pathway genes in diverse cancer type of TCGA [41]. As illustrated in Figure 5E, KISS1 was mostly negatively correlated with the immune checkpoint genes in TGCT, UCEC, STAD, COAD, BLCA and BRCA, and positively correlations predominated in THCA and PRAD. Besides, the CD274 had a relatively strong correlation with KISS1, and mainly negatively related in pan-cancer, which indicated CD274 maybe a potential checkpoint in cancers that had an abnormal KISS1 expression [42].

### Effect of KISS on treatment responses to ICIs

Based on the above results, we next focused on the potential treatment response to ICIs by KISS1. Given the importance of TMB and MSI during the decision making of an immune checkpoint therapy [43], we investigated the correlation between KISS1 expression and TMB and MSI in various cancers. As for the TMB score, the KISS1 expression was positively correlated with ESCA, STES, LUSC and negatively correlated with CESC, HNSC, THYM (Figure 6A). While, according to COAD, COADREAD, ESCA, KIRP, KIPAN, STAD, BLCA and KICH the MSI scores were positively correlated with the KISS1 expression. On the contrary, patients with PRAD, THCA and PAAD the MSI scores were negatively correlated with the KISS1 expression (Figure 6B). In terms of the tumor purity [44], which also makes sense to ICI treatment, we found that for the patients with KIRC, LIHC, READ, TGCT, BLCA, ACC, CHOL the tumor purity was positively correlated with the KISS1 expression, and the patients with PRAD showed a opposite trend (Figure 6C). In the end, we focused on the HRD [45], which is a key indicator of tumor therapy and prognosis. Interestingly, there are 9 cancers tightly correlated with KISS1, of which BRCA, KIRP, KIPAN, STAD, PRAD, HNSC, THYM, ACC have a positively relationship, and UCEC has a negatively relationship with KISS1 expression (Figure 6D).

**FIGURE 6.**
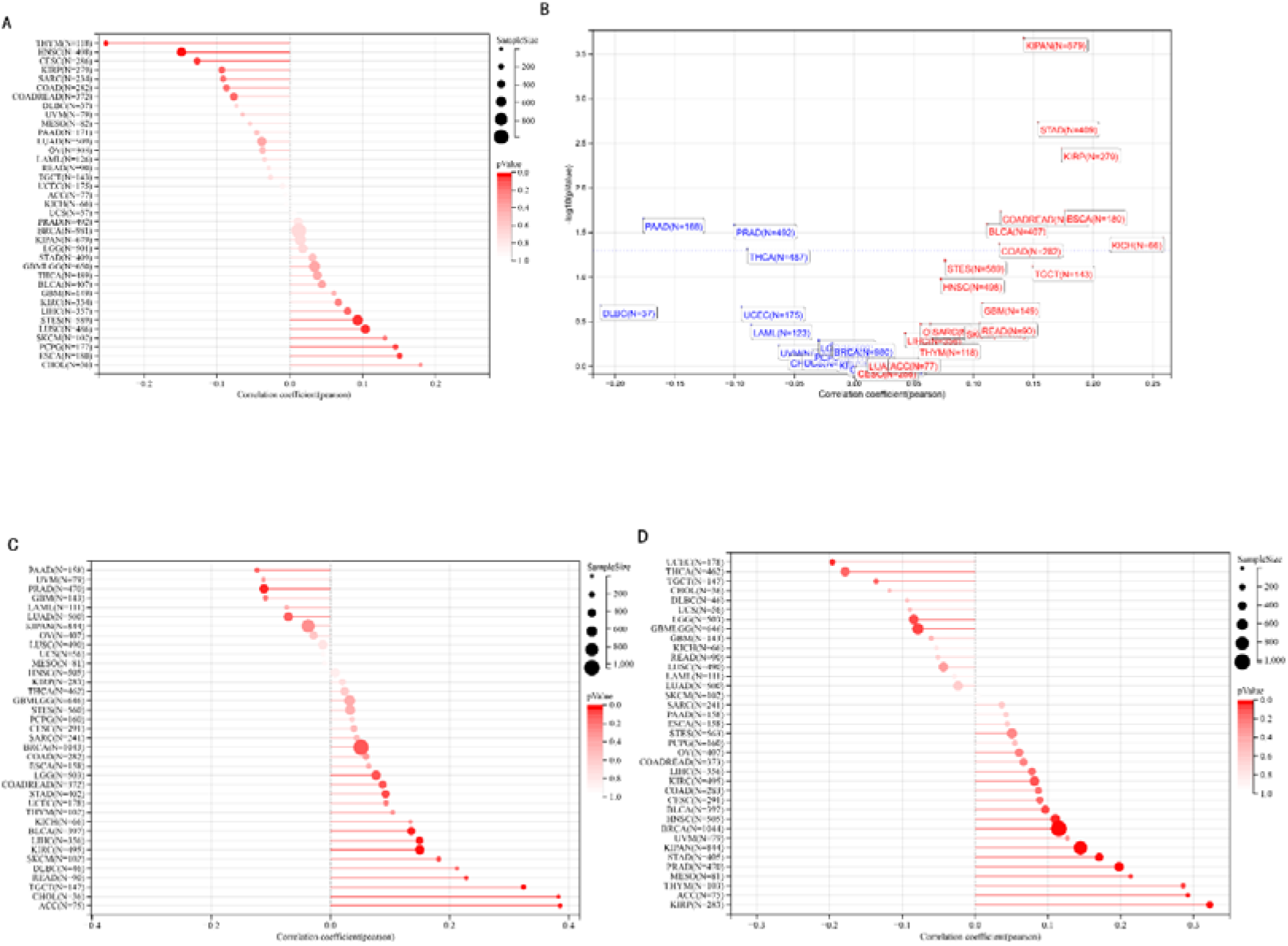
Potential effect of KISS1 on treatment responses to ICIs. (A-D) Analysis of the association between KISS1 expression and TMB, MSI, tumor purity and HRD.

### KISS1 affects breast cancer differently

From all these results, we could get a preliminary conclusion that the KISS1 affects breast cancers differently from other cancers, which is consistent with previous studies [46]. The KISS1 protein could detected in breast cancers but not in other cancers may explain why KISS1, as a metastasis suppressor, could promote the breast cancer metastasis. This raised our interesting in the potential value of KISS1 in the treatment of breast cancer. By both univariate and multivariate COX regression models, the KISS1 came out to be an independent prognostic standard, especially for different status of metastases (Figure 3C, D). Thus, it is worth to analyze the biological functions and mechanisms of KISS1 by further experiments.

### Role of KISS1 in phenotypic behavior of breast cancer cells

In order to further investigate the biological functions and mechanisms of KISS1, we decide to carry out basic experiments by breast cancer cells. Firstly, we explore the KISS1 gene expression statues in different breast cancer lines by the database from CCLE (Cancer Cell Line Encyclopedia) [47], nearly two thirds of the cell lines have a high KISS1 gene expression (Figure 7A). Then we verified the protein expression status in MDA-MB-231, which has a high capacity to develop metastasis. Indeed, the KISS1 protein expression level is higher in MDA-MB-231 than 293T (Figure 7B-7C). Importantly, the highly bone-metastatic (BM) derivatives of MDA-MB-231 had an even higher protein expression level (Figure 7B-7C). Furthermore, we generated KISS1-knockout (KISS1-KO) derivatives of MDA-MB-231 and MDA-MB-231 (BM) (Figure 7D-G). Firstly, we compared these cells for their self-renewal potential by assaying the colony formation capacity [48]. The KISS1 deficiency reduced the cell clones, and when ectopic expressing KISS1 in the KISS1-KO MDA-MB-231 the clone numbers restored (Figure 7H). And we got a similar result in bone-metastatic (BM) derivatives of MDA-MB-231 (Figure 7H). These results suggesting that KISS1 may promotes stemness of breast cancer cells in some extent. As the capacity of tumor-initiating contributes to the metastasis of a primary tumor, the clone assay may suggest the role of KISS1 in cancer metastasis [48]. Then we designed transwell assay to confirm this assumption. From Figure 7I, we could see that the migration ability was significantly reduced when the KISS1 was knocked out, whereas ectopic expression of KISS1 restored the migration ability statistically. Together, these results confirm the critical role of KISS1 in promoting metastasis in breast cancer cells.

**FIGURE 7.**
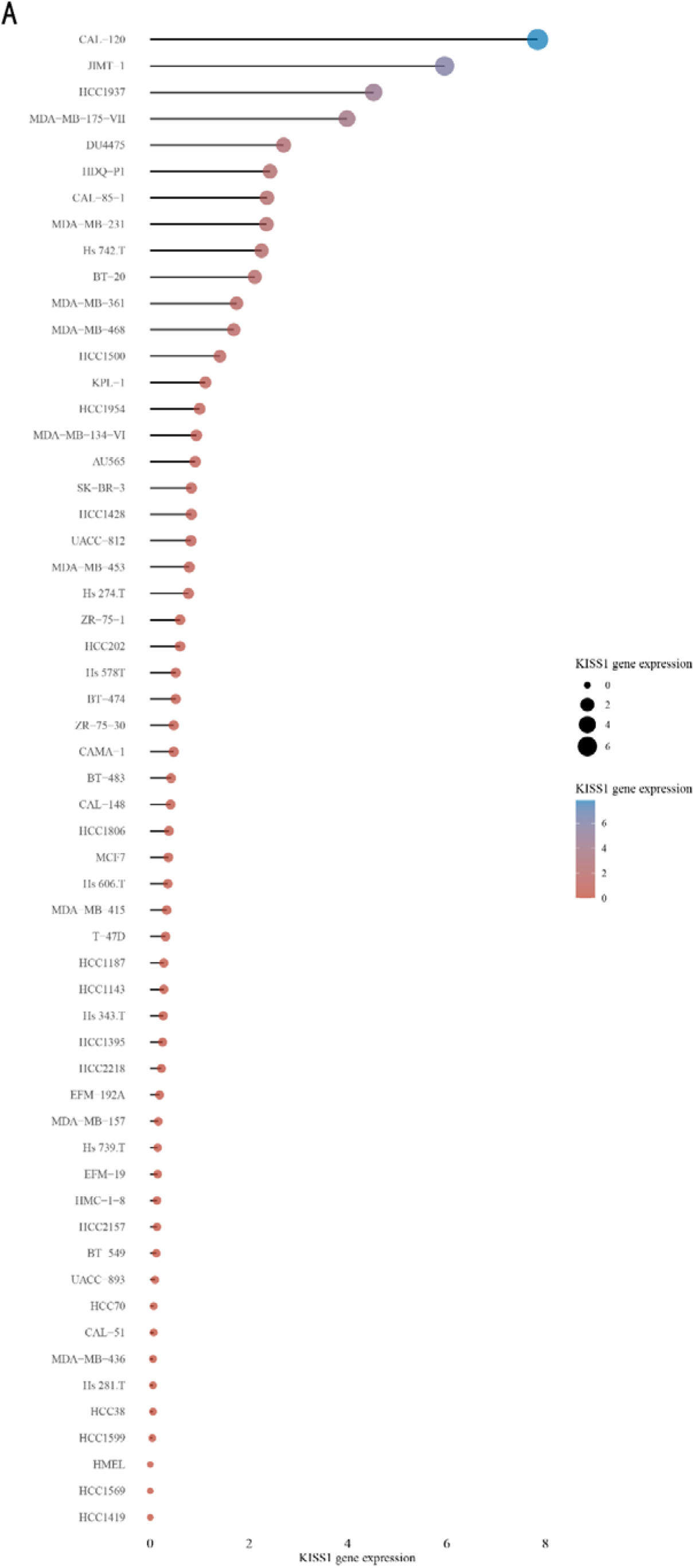
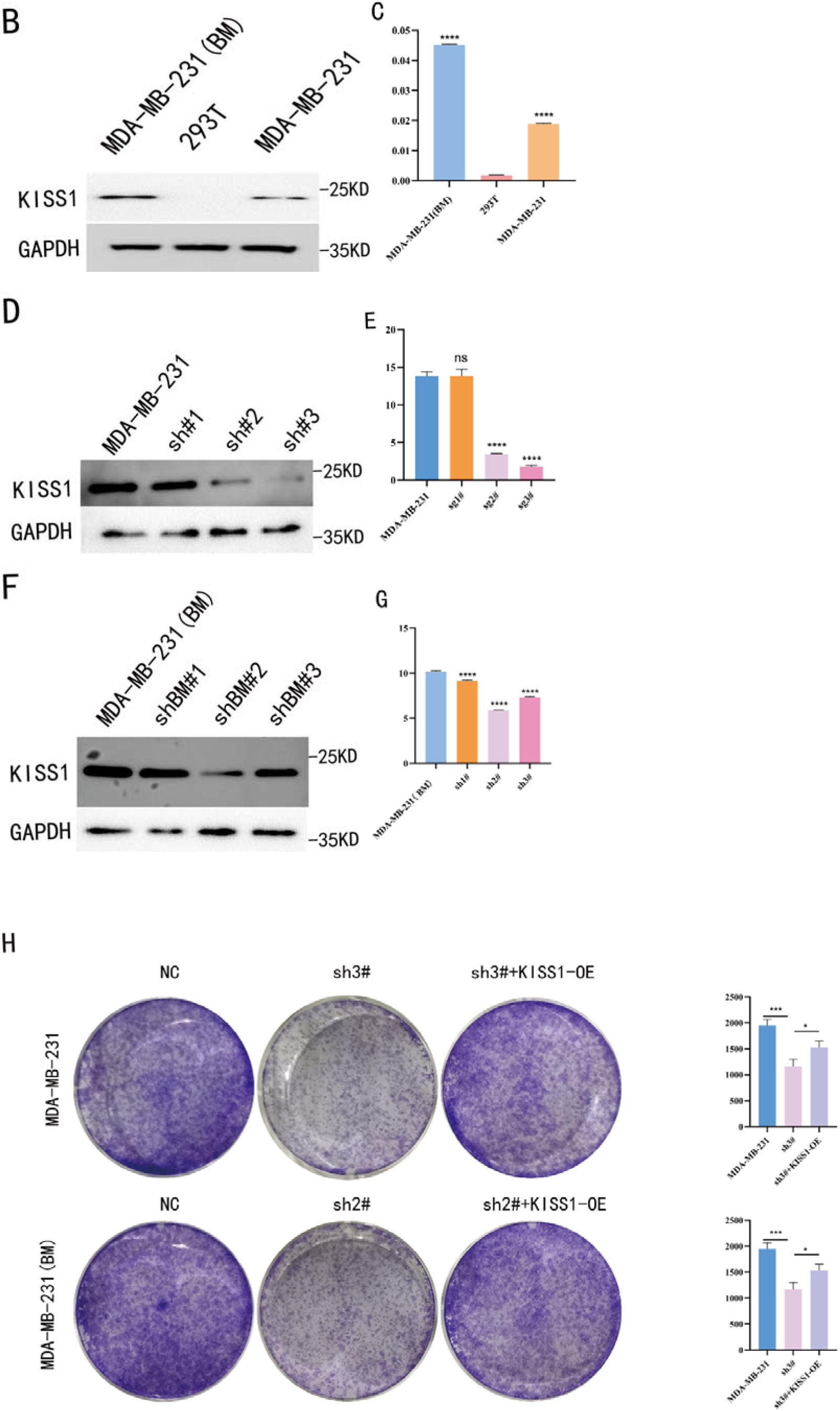
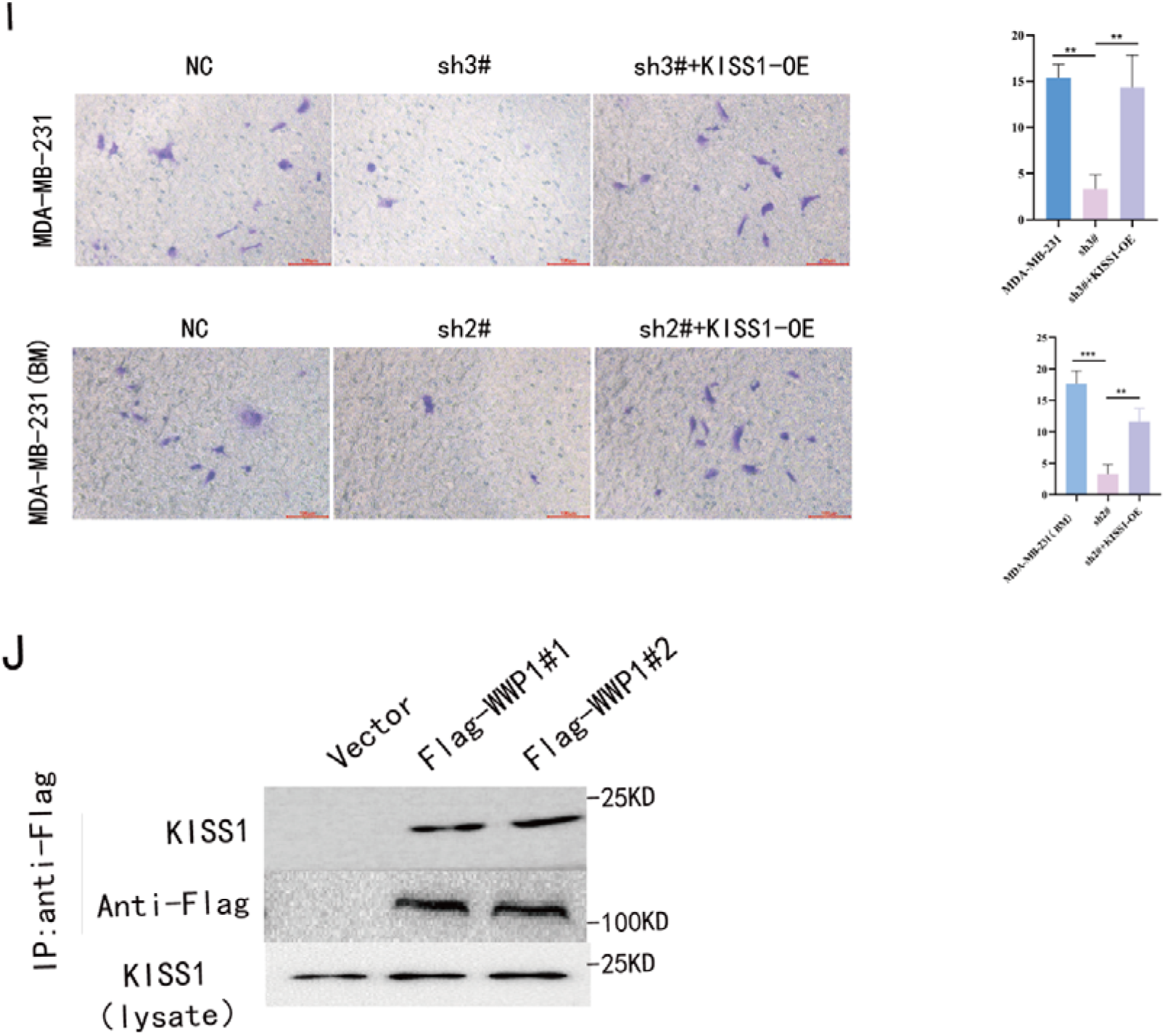
Role of KISS1 in phenotypic behavior of breast cancer cells. (A) KISS1 expression in different breast cancer cell lines. (B-G) The protein level of KISS1 in breast cancer cells and validation of KISS1 knockout. (H) Colony formation assay revels the effect of KISS1 on self-renewal potential of breast cancer cells. (I) Transwell assay detects metastasis ability. (J) WWP1 could binding to KISS1 within breast cancer cells.

### The potential Post-Translational Modification (PTMs) of KISS1 in tumors

Based on the results above, we were motivated to make it clear that why the mRNA expression level is different from the protein among the analysis from the online databases. Post-translational modification (PTMs) may play a nonnegligible role in the stability of KISS1 protein within cells [49–51]. Indeed, ubiquitination functioned dominantly in the stabilization of proteins [52]. Thus, the KISS1 may undergoing different ubiquitination in a context-dependent manner, which may explain the difference of its effect on metastasis in different cancers. To test this hypothesis, we did a binding assay to analyze whether some famous E3 ubiquitin ligases could bind with KISS1 [53–54]. Initially, we picked up WWP1 (WW domain-containing E3 ubiquitin protein ligase 1) as the candidate E3 ubiquitin ligases, for its well-known roles in regulating protein ubiquitination [55]. Interestingly, WWP1 could interacted with KISS1 within MDA-MB-231 cells (Figure 7E). Though, merely a binging assay could not confirm the ubiquitination of KISS1 within cells contributes to the influence of protein stability [53], but this indeed indicated the promising role of ubiquitination of KISS1 in cancer therapy.

## Discussion

The occurrence of metastasis events accounts for the majority of cancer-related death, and challenged the outcomes of tumor therapy [1–4]. Although, immunotherapies had reduced recurrences of various cancers, but, after a prolonged latency, still a proportion of patients developed metastases [56]. Given the low response rate of immunotherapy, more potential immunological biomarkers which could regulating tumor metastasis are needed. In one hand, KISS1 was reported as a tumor metastasis suppressor in several cancer types [12,15–17]. However, some researches find that KISS1 may contributing to the breast cancer metastasis [18]. In this study, we analyzed the physiological and pathology effect of KISS1 in pan-cancer based on the TCGA and GTEx databases. Firstly, we confirmed that the KISS1 statistically expressed highly in most cancers except KIRP KIPAN and LUSC. However, the protein expression level is not consistent to the gene expression status. It could only be detected in breast cancers tissues according to the HPA datasets. These results indicated that the KISS1 protein expressed highly in breast cancer tissues but not in other cancer tissues like Glioma, Lung cancer and Pancreatic cancer. This may partly explain the controversial role of KISS1 in regulating various cancers metastasis.

The expression difference of KISS1 among various cancers may suggest the biomarker potential of KISS1 and we further tested this potential in background of clinical and metastatic stage. For the same tumor, the KISS1 expressed differently among varies pathological stages. Besides, in terms of different metastasis status, the difference of KISS1 expression only make sense in COADREAD, KIPAN, HNSC, PAAD, SKCM. These results, based on the gene expression database from TGCA, was contrast to some basic researches [12,15–18], which way partly be explained by the occurrence of modifications of KISS1 protein in cancer cells after translation [49–51].

In this study, it is the first time to analyzing the prospective role of KISS1 in immunotherapy in pan-cancer. Began with the relationship between KISS1 expression and immune cell infiltration, we adopted three immune scores, including StromalScore, ImmuneScore and EstimateScore [12]. In summary, KISS1 correlated tightly with all the three scores, and mostly the correlation was negative. For example, in BRCA the immune cell infiltration level is lower while there is a higher expression level of KISS1 according to the three immune scores. However, this may partly explain the crucial role of KISS1 in cancers including breast cancer. Accordingly, the conclusions presented by three scores might give us a conceptual map of immune and stromal cell in various cancers which may sort of explaining the differential role of KISS1 in the metastasis of malignancies [33]. Based on this, we next targeting suppressive tumor microenvironment (STM), as the pre-existence of immunosuppressive cells is a major reason for the failure of immunotherapy for tumors [57]. We analyzed the potential immunomodulatory effects of KISS1 by evaluating its efficacy on Myeloid-derived suppressor cells (MDSCs) and regulatory T cells (Tregs) [58–59]. Comparations between these results suggested that the immunosuppressive cells infiltrate highly in BRCA, BRCA-LumA, KIRC, KIRP and MESO, where there is a high KISS1 expression too. Take BRCA, especially LumA subtype, for example, the infiltrations of MDSC and Tregs are positively correlated with KISS1 expression according to the TIED, CIBERSORT and CIBERSORT-ABS algorithms. Assumingly, this may have some relationship with the breast cancer metastasis, especially the bone preference [5]. Further investigations into these mechanisms might help to explain the higher frequency of bone metastasis of luminal-like subtype breast cancers. Thus, novel effective immunotherapies exclusive for cancer metastasis may come out [57]. In another hand, as the infiltration of immunosuppressive cells may compromising the therapeutic efficacy of ICIs, the study was followed by the analysis of the correlation of KISS1 expression with ICI-related genes [60]. A significant correlation was found, which dominated by positive correlation. Interestingly, CD274 (PD-L1) showed a relatively strong correlation with KISS1. Concordantly, the PD-L1 plays an important role in the immunotherapy of breast cancers [61], which reinforced the potential role of KISS1 in the treatment of breast cancer.

Given the unique role of KISS1 in breast cancer, we further analyzed that the KISS1 may be an independent predictor of tumor metastases in breast cancer patients. Then we tested the possibility that knocking down of the KISS1 gene could statistically reduce the metastasis capacity of MBA-MD-231 cells, which implied the EMT process of tumor cells was influenced, to some extent, by the KISS1.

In summary, our first pan-cancer analysis and experiments of KISS1, once reported as a metastasis suppressor that expressed differently in various tumors, implied statistical correlation of KISS1 expression with clinical prognosis, immune cell infiltration, including immunosuppressive cells and lymphocytes, and tumor metastasis capacity, which add to the understanding of the epigenetic regulatory mechanism in terms of clinical tumor samples. Indeed, our study confirm the therapeutic and prognostic marker potential of KISS1 and indicate the potential of immunotherapy target for tumor metastasis. However, the lacking of in-depth clinical and basic experiments restrict the verification of its role as an immunotherapeutic target, and calling for further in-depth studies.

